# The creation-mutation-selection model: simulation and the advantage of sex

**DOI:** 10.1101/2023.08.12.553090

**Authors:** Gordon Irlam

## Abstract

The creation-mutation-selection model makes predictions regarding the fitness of asexual and sexual populations in an environment that includes both positive and negative selection. However, for the asexual case, the model predictions depend upon the variance or skewness of the population log fitness distribution, something which the model does not provide. This makes the analytical comparison of the fitness of asexual and sexual populations appear to be intractable. Instead, simulation is required. The fitness of asexual and sexual populations is simulated over a range of plausible parameter values. The fitness is found to agree with the predictions of the creation-mutation-selection model. The cost of asexuality is found to exceed the cost of sex by a very large margin for eukaryotic species today, and probably for the first eukaryotes. Prokaryotes and the mitochondrion are viewed as asexual species for which asexuality incurs little cost.

## Introduction

The creation-mutation-selection model is a mathematical model for the fitness of asexual and sexual populations in the presence of positive and negative selection[1]. Unfortunately, for an asexual population, the model only predicts the fitness of the population if the variance or non-standardized skewness of the population log fitness distribution is known. Thus to compare the fitness of asexual and sexual populations it is necessary to resort to simulation.

Computer simulations of the asexual and sexual models was performed, and compared to the predictions of the creation-mutation-selection model. The model depends on several parameters. Reasonable values for these parameters have been estimated previously, and were used for the simulations[2].

Reviewing the creation-mutation-selection model, consider a population of *N* haploid organisms in a Wright-Fisher like discrete time model of non-overlapping generations. Suppose positive mutational opportunities occur with mean rate Γ_*p*_ per generation across all organisms, and negative mutational opportunities occur with mean rate Γ_*n*_ per organism. Let *μ*_*ss*_ be the spontaneous mutation rate, that is the rate of satisfaction of mutational opportunity sites per sexual or asexual generation. Let *s* be the selection coefficient associated with the satisfaction of a mutational opportunity site. For sexual organisms selection is assumed to occur in haploids, or equivalently in diploids with a selection coefficient of roughly 2*s* and a heterozygous effect of 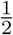. When 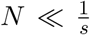 the sites are termed neutral sites, and the positive or negative site creation rate is denoted Γ_0_. An organism with no mutational opportunity sites has a fitness of 1. Fitness when multiple sites are present is assumed to combine additively in log space, and log fitness will be a negative value. Natural selection involves selection with replacement in accordance with fitness. Sex involves drawing a new population of organisms by mating two randomly selected organisms at a time, and randomly selecting mutational opportunities from each of them without linkage.

For presently existing eukaryotes, genomic analysis suggests Γ_*p*_ is typically in the range 10^*−*3^ to 10^*−*2^ population wide adaptive mutational opportunities per sexual generation[2]. Smaller and larger vales for Γ_*p*_ are considered, partially to gain understanding, and partially to speculate that prokaryotes experience smaller rates of positive selection. The range for Γ_*p*_ of 10^*−*4^ to 10^*−*1^ will be considered.

For eukaryotes, Γ_*n*_ is typically in the range 10^*−*1^ to 10^1^ negative sites per organism per sexual generation[2]. Values two orders of magnitude smaller than this are also considered so as to start to encompass the range spanned by prokaryotes. Prokaryotes have spontaneous mutation rates per genome in the range 10^*−*4^ to 10^*−*2^ per generation[3][Figure 1, visual inspection assuming all sites are under negative selection]. The range for Γ_*n*_ of 10^*−*3^ to 10^1^ will be considered.

**Figure 1:**
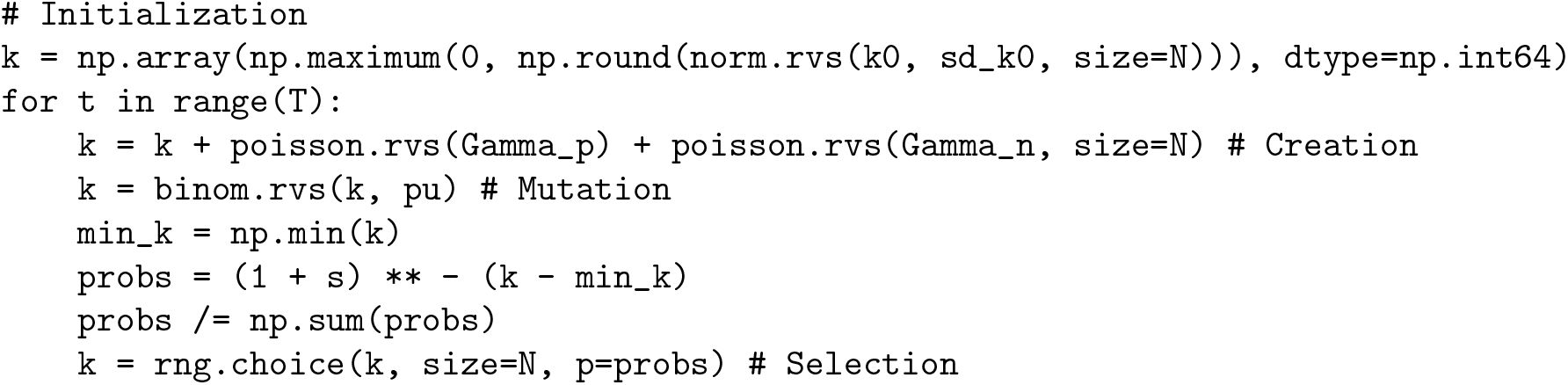
NumPy Python pseudo-code for the asexual simulation.

For eukaryotes, *μ*_*ss*_ is typically in the range 10^*−*9^ to 10^*−*8^ spontaneous mutations per site per sexual generation[2]. For bacteria the spontaneous mutation rate per base pair, *μ*_*bs*_, ranges from 1.4*×*10^*−*10^ to 4*×*10^*−*9^, placing *μ*_*ss*_ roughly in the range 10^*−*10^ to 10^*−*9^[3][Supplement 1]. Rates for archaea are even smaller, at around 10^*−*11^ per generation[3][Supplement 1]. The range for *μ*_*ss*_ of 10^*−*10^ to 10^*−*8^ will be considered.

Let *x* be the log fitness of an organism, and let 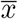 be the mean log fitness of the population. So they may be referred to in this paper, in the creation-mutation-selection model the key formulas for 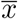 are as given below[1].

For asexual organisms, fitness expressed in terms of 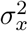, the variance of *x*,

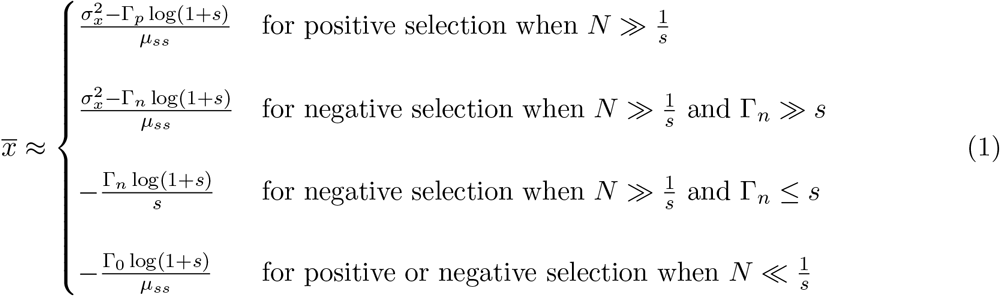

For asexual organisms, fitness expressed in terms of *μ*_3,*x*_, the third central moment of *x*, that is non-standardized skewness,

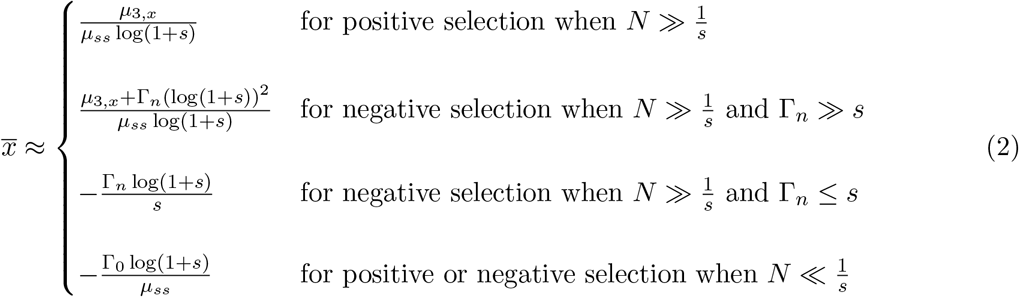

For sexual organisms:

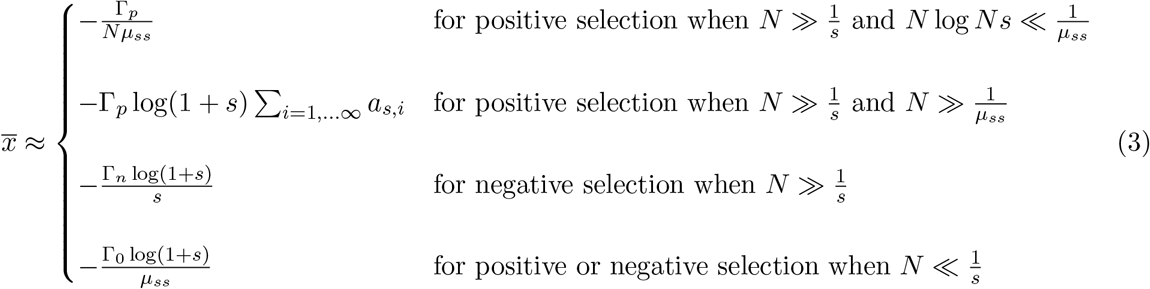

where

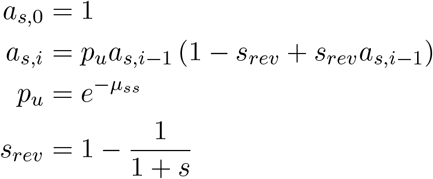

For positive and negative selection, or for multiple selection coefficients, log fitness values may be added in the case of sexual populations. For asexual populations the situation is more complex.

As will be shown, the results of computer simulation agree with the mathematical creation-mutation-selection model, thus validating the model.

Having computed the fitness in the asexual case it becomes possible to determine the advantage of sex. This is done for prokaryotes, present day eukaryotes, the first eukaryotes, and the asexual mitochondrion. Sex appears to have an advantage where it occurs, and not to have an advantage where it doesn’t.

## Results

The selection coefficients for sites as a result of negative selection are likely to be larger than for positive selection. There are many ways a protein could mutate under negative selection rendering it almost non-functional, but the likelihood of a positive mutation of similar magnitude is likely to be rare.

### Results for the asexual model

A computer simulation of the asexual case was developed. A default population size of 10^5^ is used. This is for ease of simulation. Values ranging from 10^4^ to 10^8^ are also considered. Values any larger than this would require an excessive amount of simulation time.

### Positive selection and asexual mean log fitness

The observed mean log fitness of the asexual model under positive selection is shown in Table 1. It should be compared to Table 2 which shows the predicted log fitness calculated based on the mean non-standardized skewness of the log fitness distribution (using equation 2). We could have equally well have chosen to predict log fitness based on the mean variance of the log fitness distribution (using equation 1). To estimate the average actual fitness values raise *e* to the power of these mean log fitness values.

**Table 1:** Observed mean log fitness for the asexual case under positive selection. Log fitness values are relative to a perfectly adapted organism having a log fitness of zero. *μ*_*ss*_ is the rate of mutation per site per generation. *N* is the population size. Γ_*p*_ is the rate of positive site creation per generation. Γ_*n*_ is the rate of deleterious site creation per organism per generation. *s* is the constant selection coefficient for each site per generation.

| $\mu_{ss}$ | $N$ | $\Gamma_p$ | $10^{-1}$ | $10^{-2}$ | $10^{-3}$ | $10^{-4}$ |
| --- | --- | --- | --- | --- | --- | --- |
| $10^{-8}$ | $10^5$ | $10^{-4}$ | -0.0557 | -0.0483 | -0.0883 | -0.169 |
| $10^{-8}$ | $10^5$ | $10^{-3}$ | -0.596 | -1.36 | -7.45 | -3.85 |
| $10^{-8}$ | $10^5$ | $10^{-2}$ | -30.5 | -387 | -284 | -60.9 |
| $10^{-8}$ | $10^5$ | $10^{-1}$ | $-2.34 \times 10^4$ | $-2.24 \times 10^4$ | $-5.45 \times 10^3$ | -782 |
| $10^{-9}$ | $10^8$ | $10^{-2}$ | -6.99 | -862 | $-1.5 \times 10^3$ | -464 |
| $10^{-9}$ | $10^7$ | $10^{-2}$ | -20.4 | $-1.37 \times 10^3$ | $-1.78 \times 10^3$ | -502 |
| $10^{-9}$ | $10^6$ | $10^{-2}$ | -72.4 | $-2.18 \times 10^3$ | $-2.19 \times 10^3$ | -540 |
| $10^{-9}$ | $10^5$ | $10^{-4}$ | -0.504 | -0.496 | -0.878 | -1.65 |
| $10^{-9}$ | $10^5$ | $10^{-3}$ | -5.8 | -14.2 | -75.6 | -37.6 |
| $10^{-9}$ | $10^5$ | $10^{-2}$ | -309 | $-3.85 \times 10^3$ | $-2.84 \times 10^3$ | -614 |
| $10^{-9}$ | $10^5$ | $10^{-1}$ | $-2.34 \times 10^5$ | $-2.24 \times 10^5$ | $-5.46 \times 10^4$ | $-7.82 \times 10^3$ |
| $10^{-9}$ | $10^4$ | $10^{-2}$ | $-1.57 \times 10^3$ | $-7.9 \times 10^3$ | $-3.94 \times 10^3$ | -755 |
| $10^{-10}$ | $10^5$ | $10^{-4}$ | -4.71 | -4.77 | -8.84 | -16.2 |
| $10^{-10}$ | $10^5$ | $10^{-3}$ | -55.7 | -143 | -732 | -385 |
| $10^{-10}$ | $10^5$ | $10^{-2}$ | $-2.76 \times 10^3$ | $-3.9 \times 10^4$ | $-2.82 \times 10^4$ | $-6.19 \times 10^3$ |
| $10^{-10}$ | $10^5$ | $10^{-1}$ | $-2.37 \times 10^6$ | $-2.23 \times 10^6$ | $-5.46 \times 10^5$ | $-7.82 \times 10^4$ |

**Table 2:**
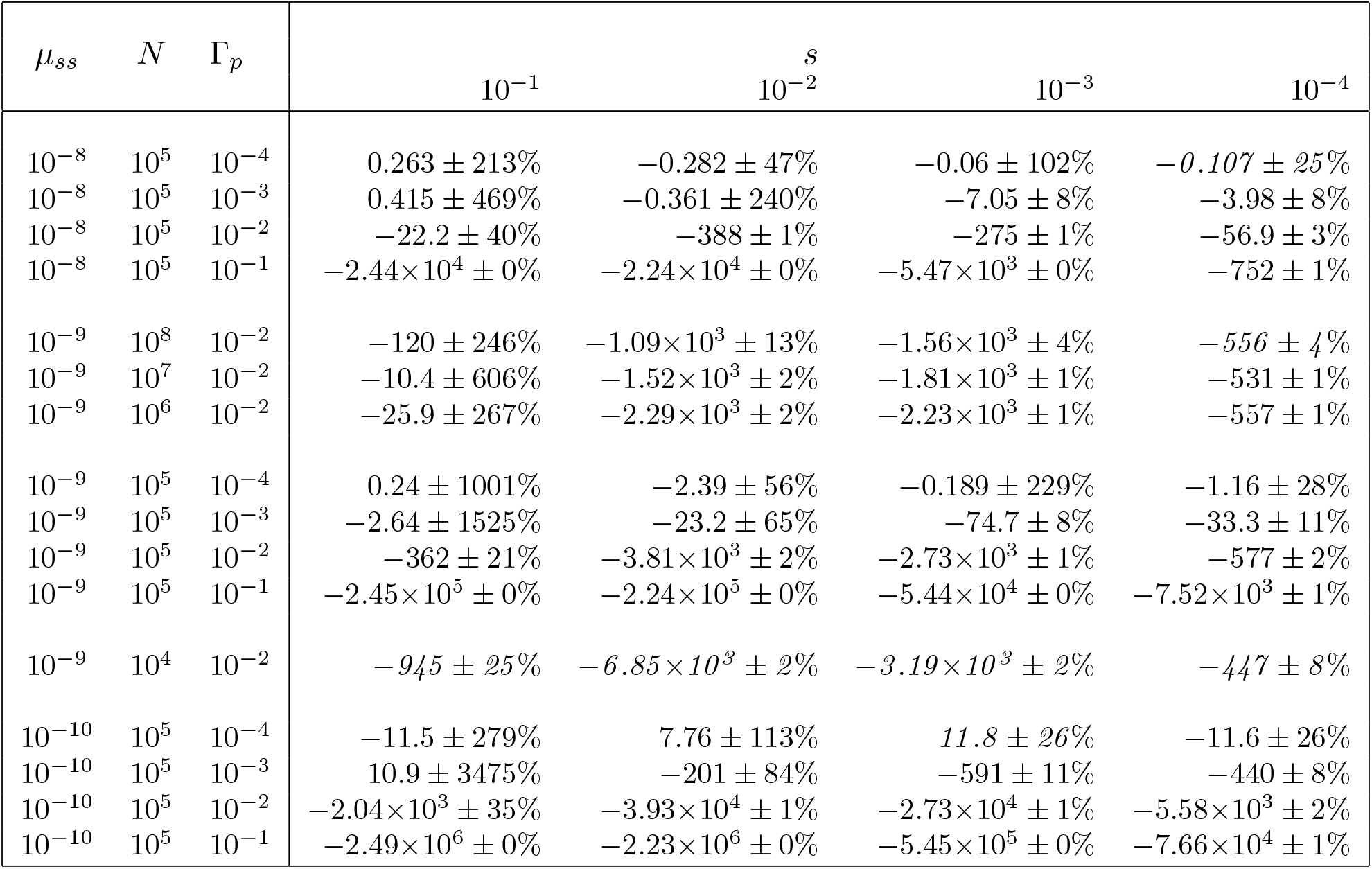
Predicted mean log fitness and standard error for the asexual case under positive selection based on the observed log fitness non-standardized skewness. Predictions whose confidence intervals failed to overlap the observed values are indicated by italics. Symbols are as described in Table 1.

| $\mu_{ss}$ | $N$ | $\Gamma_p$ | $10^{-1}$ | $10^{-2}$ | $10^{-3}$ | $10^{-4}$ |
| --- | --- | --- | --- | --- | --- | --- |
| $10^{-8}$ | $10^5$ | $10^{-4}$ | $0.263 \pm 213\%$ | $-0.282 \pm 47\%$ | $-0.06 \pm 102\%$ | $-0.107 \pm 25\%$ |
| $10^{-8}$ | $10^5$ | $10^{-3}$ | $0.415 \pm 469\%$ | $-0.361 \pm 240\%$ | $-7.05 \pm 8\%$ | $-3.98 \pm 8\%$ |
| $10^{-8}$ | $10^5$ | $10^{-2}$ | $-22.2 \pm 40\%$ | $-388 \pm 1\%$ | $-275 \pm 1\%$ | $-56.9 \pm 3\%$ |
| $10^{-8}$ | $10^5$ | $10^{-1}$ | $-2.44 \times 10^4 \pm 0\%$ | $-2.24 \times 10^4 \pm 0\%$ | $-5.47 \times 10^3 \pm 0\%$ | $-752 \pm 1\%$ |
| $10^{-9}$ | $10^8$ | $10^{-2}$ | $-120 \pm 246\%$ | $-1.09 \times 10^3 \pm 13\%$ | $-1.56 \times 10^3 \pm 4\%$ | $-556 \pm 4\%$ |
| $10^{-9}$ | $10^7$ | $10^{-2}$ | $-10.4 \pm 606\%$ | $-1.52 \times 10^3 \pm 2\%$ | $-1.81 \times 10^3 \pm 1\%$ | $-531 \pm 1\%$ |
| $10^{-9}$ | $10^6$ | $10^{-2}$ | $-25.9 \pm 267\%$ | $-2.29 \times 10^3 \pm 2\%$ | $-2.23 \times 10^3 \pm 1\%$ | $-557 \pm 1\%$ |
| $10^{-9}$ | $10^5$ | $10^{-4}$ | $0.24 \pm 1001\%$ | $-2.39 \pm 56\%$ | $-0.189 \pm 229\%$ | $-1.16 \pm 28\%$ |
| $10^{-9}$ | $10^5$ | $10^{-3}$ | $-2.64 \pm 1525\%$ | $-23.2 \pm 65\%$ | $-74.7 \pm 8\%$ | $-33.3 \pm 11\%$ |
| $10^{-9}$ | $10^5$ | $10^{-2}$ | $-362 \pm 21\%$ | $-3.81 \times 10^3 \pm 2\%$ | $-2.73 \times 10^3 \pm 1\%$ | $-577 \pm 2\%$ |
| $10^{-9}$ | $10^5$ | $10^{-1}$ | $-2.45 \times 10^5 \pm 0\%$ | $-2.24 \times 10^5 \pm 0\%$ | $-5.44 \times 10^4 \pm 0\%$ | $-7.52 \times 10^3 \pm 1\%$ |
| $10^{-9}$ | $10^4$ | $10^{-2}$ | $-945 \pm 25\%$ | $-6.85 \times 10^3 \pm 2\%$ | $-3.19 \times 10^3 \pm 2\%$ | $-447 \pm 8\%$ |
| $10^{-10}$ | $10^5$ | $10^{-4}$ | $-11.5 \pm 279\%$ | $7.76 \pm 113\%$ | $11.8 \pm 26\%$ | $-11.6 \pm 26\%$ |
| $10^{-10}$ | $10^5$ | $10^{-3}$ | $10.9 \pm 3475\%$ | $-201 \pm 84\%$ | $-591 \pm 11\%$ | $-440 \pm 8\%$ |
| $10^{-10}$ | $10^5$ | $10^{-2}$ | $-2.04 \times 10^3 \pm 35\%$ | $-3.93 \times 10^4 \pm 1\%$ | $-2.73 \times 10^4 \pm 1\%$ | $-5.58 \times 10^3 \pm 2\%$ |
| $10^{-10}$ | $10^5$ | $10^{-1}$ | $-2.49 \times 10^6 \pm 0\%$ | $-2.23 \times 10^6 \pm 0\%$ | $-5.45 \times 10^5 \pm 0\%$ | $-7.66 \times 10^4 \pm 1\%$ |

While correct according to our mathematical model, it should be noted that a few of the values reported in Table 1 are so large as to be biologically invalid. E.g. for *μ*_*ss*_ = 10^*−*10^, *N* = 10^5^, Γ_*p*_ = 10^*−*1^, and *s* = 10^*−*4^ we have an observed mean log fitness, 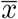, of *−*7.82*×*10^4^. Such a fitness, requires there be an average of 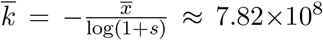 unsatisfied sites per organism. This exceeds the length of the region under the control of negative selection for many eukaryotic genomes. The large values are just the model’s way of expressing the infeasibility of an asexual organism existing with these parameter values.

As *N* increases, mean log fitness becomes smaller. However the rate at which it shrinks is not excessive. This means that most of the conclusions to be reached based on smaller population sizes should still be valid for large populations, such as 10^9^ or more, which were not simulated.

The reported values for mean log fitness, 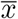, have a standard error of 2.5% or less, apart from seventeen small values of 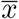 all of which were computed by direct roll-out and for which standard error values are variable. Predictions for 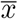 include standard error estimates indicated by the plus/minus symbol based on the observed variability of *μ*_3,*x*_. Predictions whose plus or minus two standard error confidence interval fails to overlap a plus or minus 5% confidence interval for 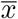 are indicated by italics. Under the hypothesis that the mathematical model is accurate this should happen less than 5% of the time.

Despite typically running for 1,000,000 generations the mean value of *μ*_3,*x*_ often involves considerable standard error. Referring to equation 2, when *μ*_3,*x*_ is small this directly produces a large percentage standard error in 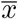. If expressed in absolute terms, the standard error would be no larger for small values of 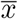 than for large values.

Predictions based on *μ*_3,*x*_ correctly predicted the value of 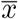 except primarily when *N* = 10^4^. It is reasonable to assume that the discrete nature of small populations causes a loss in fidelity of *μ*_3,*x*_, causing predictions based on it to be inaccurate.

### Negative selection and asexual mean log fitness

The observed mean log fitness of the asexual model under negative selection is shown in Table 3. It should be compared to Table 4 which shows the predicted log fitness calculated based on the mean non-standardized skewness or the rate of deleterious site creation (using equation 2).

**Table 3:** Observed mean log fitness for the asexual case under negative selection. Symbols are as described in Table 1.

| $\mu_{ss}$ | $N$ | $\Gamma_n$ | $10^{-1}$ | $10^{-2}$ | $10^{-3}$ | $10^{-4}$ |
| --- | --- | --- | --- | --- | --- | --- |
| $10^{-8}$ | $10^5$ | $10^{-3}$ | -0.000953 | -0.000995 | -0.00101 | -1.14 |
| $10^{-8}$ | $10^5$ | $10^{-2}$ | -0.00953 | -0.00996 | -50.8 | -41.2 |
| $10^{-8}$ | $10^5$ | $10^{-1}$ | -0.0953 | $-1.79 \times 10^3$ | $-3.23 \times 10^3$ | -659 |
| $10^{-8}$ | $10^5$ | 1 | $-7.67 \times 10^4$ | $-2.67 \times 10^5$ | $-6.06 \times 10^4$ | $-8.19 \times 10^3$ |
| $10^{-8}$ | $10^5$ | $10^1$ | $-2.46 \times 10^7$ | $-5.7 \times 10^6$ | $-7.92 \times 10^5$ | $-9.08 \times 10^4$ |
| $10^{-9}$ | $10^8$ | 1 | -0.953 | $-1.61 \times 10^6$ | $-5.04 \times 10^5$ | $-7.53 \times 10^4$ |
| $10^{-9}$ | $10^7$ | 1 | -12.4 | $-1.88 \times 10^6$ | $-5.32 \times 10^5$ | $-7.7 \times 10^4$ |
| $10^{-9}$ | $10^6$ | 1 | $-1.28 \times 10^5$ | $-2.22 \times 10^6$ | $-5.65 \times 10^5$ | $-7.93 \times 10^4$ |
| $10^{-9}$ | $10^5$ | $10^{-3}$ | -0.000953 | -0.0762 | -0.00101 | -11.4 |
| $10^{-9}$ | $10^5$ | $10^{-2}$ | -0.00953 | -0.00996 | -504 | -413 |
| $10^{-9}$ | $10^5$ | $10^{-1}$ | -0.0953 | $-1.78 \times 10^4$ | $-3.24 \times 10^4$ | $-6.62 \times 10^3$ |
| $10^{-9}$ | $10^5$ | 1 | $-7.65 \times 10^5$ | $-2.67 \times 10^6$ | $-6.06 \times 10^5$ | $-8.19 \times 10^4$ |
| $10^{-9}$ | $10^5$ | $10^1$ | $-2.46 \times 10^8$ | $-5.7 \times 10^7$ | $-7.92 \times 10^6$ | $-9.08 \times 10^5$ |
| $10^{-9}$ | $10^4$ | 1 | $-2.46 \times 10^6$ | $-3.29 \times 10^6$ | $-6.63 \times 10^5$ | $-8.58 \times 10^4$ |
| $10^{-10}$ | $10^5$ | $10^{-3}$ | -0.5 | -0.000995 | -0.00101 | -113 |
| $10^{-10}$ | $10^5$ | $10^{-2}$ | -0.00953 | -0.00996 | $-5.25 \times 10^3$ | $-4.14 \times 10^3$ |
| $10^{-10}$ | $10^5$ | $10^{-1}$ | -0.0953 | $-1.8 \times 10^5$ | $-3.24 \times 10^5$ | $-6.55 \times 10^4$ |
| $10^{-10}$ | $10^5$ | 1 | $-7.65 \times 10^6$ | $-2.67 \times 10^7$ | $-6.06 \times 10^6$ | $-8.19 \times 10^5$ |
| $10^{-10}$ | $10^5$ | $10^1$ | $-2.46 \times 10^9$ | $-5.7 \times 10^8$ | $-7.92 \times 10^7$ | $-9.08 \times 10^6$ |

**Table 4:** Predicted mean log fitness and standard error for the asexual case under negative selection based on the observed log fitness non-standardized skewness or the rate of deleterious site creation. Symbols are as described in Table 1.

| $\mu_{ss}$ | $N$ | $\Gamma_n$ | $s$ | | | |
| --- | --- | --- | --- | --- | --- | --- |
| | | | $10^{-1}$ | $10^{-2}$ | $10^{-3}$ | $10^{-4}$ |
| $10^{-8}$ | $10^5$ | $10^{-3}$ | -0.000953 | -0.000995 | -0.001 | $-2.19 \pm 22\%$ |
| $10^{-8}$ | $10^5$ | $10^{-2}$ | -0.00953 | -0.00995 | $-45 \pm 8\%$ | $-35.2 \pm 7\%$ |
| $10^{-8}$ | $10^5$ | $10^{-1}$ | -0.0953 | $-1.19 \times 10^3 \pm 3\%$ | $-3.17 \times 10^3 \pm 0\%$ | $-637 \pm 2\%$ |
| $10^{-8}$ | $10^5$ | 1 | $3.68 \times 10^5 \pm 0\%$ | $-2.62 \times 10^5 \pm 0\%$ | $-6.01 \times 10^4 \pm 0\%$ | $-8.1 \times 10^3 \pm 1\%$ |
| $10^{-8}$ | $10^5$ | $10^1$ | $-2.13 \times 10^7 \pm 0\%$ | $-5.67 \times 10^6 \pm 0\%$ | $-7.89 \times 10^5 \pm 0\%$ | $-8.98 \times 10^4 \pm 0\%$ |
| $10^{-9}$ | $10^8$ | 1 | $4.47 \times 10^6 \pm 0\%$ | $-1.57 \times 10^6 \pm 0\%$ | $-5.09 \times 10^5 \pm 0\%$ | $-7.63 \times 10^4 \pm 1\%$ |
| $10^{-9}$ | $10^7$ | 1 | $4.47 \times 10^6 \pm 0\%$ | $-1.84 \times 10^6 \pm 0\%$ | $-5.34 \times 10^5 \pm 0\%$ | $-7.89 \times 10^4 \pm 0\%$ |
| $10^{-9}$ | $10^6$ | 1 | $4.34 \times 10^6 \pm 0\%$ | $-2.19 \times 10^6 \pm 0\%$ | $-5.66 \times 10^5 \pm 0\%$ | $-8.07 \times 10^4 \pm 0\%$ |
| $10^{-9}$ | $10^5$ | $10^{-3}$ | -0.000953 | $-0.000995$ | -0.001 | $-30.8 \pm 20\%$ |
| $10^{-9}$ | $10^5$ | $10^{-2}$ | -0.00953 | -0.00995 | $-426 \pm 10\%$ | $-352 \pm 7\%$ |
| $10^{-9}$ | $10^5$ | $10^{-1}$ | -0.0953 | $-1.1 \times 10^4 \pm 4\%$ | $-3.15 \times 10^4 \pm 0\%$ | $-6.29 \times 10^3 \pm 2\%$ |
| $10^{-9}$ | $10^5$ | 1 | $3.69 \times 10^6 \pm 0\%$ | $-2.62 \times 10^6 \pm 0\%$ | $-6.02 \times 10^5 \pm 0\%$ | $-8.08 \times 10^4 \pm 1\%$ |
| $10^{-9}$ | $10^5$ | $10^1$ | $-2.13 \times 10^8 \pm 0\%$ | $-5.67 \times 10^7 \pm 0\%$ | $-7.89 \times 10^6 \pm 0\%$ | $-8.99 \times 10^5 \pm 0\%$ |
| $10^{-9}$ | $10^4$ | 1 | $2.01 \times 10^6 \pm 0\%$ | $-3.17 \times 10^6 \pm 0\%$ | $-6.21 \times 10^5 \pm 0\%$ | $-6.84 \times 10^4 \pm 2\%$ |
| $10^{-10}$ | $10^5$ | $10^{-3}$ | $-0.000953$ | -0.000995 | -0.001 | $-200 \pm 23\%$ |
| $10^{-10}$ | $10^5$ | $10^{-2}$ | -0.00953 | -0.00995 | $-5.41 \times 10^3 \pm 9\%$ | $-3.52 \times 10^3 \pm 8\%$ |
| $10^{-10}$ | $10^5$ | $10^{-1}$ | -0.0953 | $-1.21 \times 10^5 \pm 3\%$ | $-3.2 \times 10^5 \pm 0\%$ | $-6.37 \times 10^4 \pm 2\%$ |
| $10^{-10}$ | $10^5$ | 1 | $3.69 \times 10^7 \pm 0\%$ | $-2.63 \times 10^7 \pm 0\%$ | $-6.01 \times 10^6 \pm 0\%$ | $-8.06 \times 10^5 \pm 1\%$ |
| $10^{-10}$ | $10^5$ | $10^1$ | $-2.13 \times 10^9 \pm 0\%$ | $-5.67 \times 10^8 \pm 0\%$ | $-7.89 \times 10^7 \pm 0\%$ | $-8.99 \times 10^6 \pm 0\%$ |

For the most part, predictions based on negative skewness or the rate of deleterious site creation match the observed values. The predictions for Γ_*n*_ = 10^*−*1^ and *s* = 10^*−*2^, and Γ_*n*_ *≥* 1 and *s* = 10^*−*1^ do not match. This appears to be the increasingly broad transition region from the existence of zero or one mutational opportunity site to a continuous fitness distribution. Two of the measured values for Γ_*n*_ = 10^*−*3^, namely *μ*_*ss*_ = 10^*−*9^ and *s* = 10^*−*2^, and *μ*_*ss*_ = 10^*−*10^ and *s* = 10^*−*1^ are small as predicted, but still anomalous, possibly because the simulation wasn’t run for long enough in these cases. Running the first of these scenarios a second time with a different random seed produced -0.000997, a value in line with predictions. The originally reported values are used in the subsequent analysis.

### Results for the sexual model

A computer simulation of the sexual model was developed. Simulating large values of *N* has a high computational cost, so the simulation was restricted to values equal to or smaller than 10^6^.

### Positive selection and sexual mean log fitness

For positive selection we are interested in assessing the accuracy of both the queuing formula and the recurrence formula. However the recurrence formula requires that 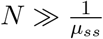, which is larger than the *N* values which can reasonably be simulated. Consequently to aid in verifying the recurrence formula we also use a non-physical *μ*_*ss*_ value of 10^*−*4^.

Tables 5 show the observed mean log fitness for the simulated sexual model. Table 6 shows mean log fitness calculated using the larger of the queuing formula or recurrence formula (equation 3). This means for *μ*_*ss*_ = 10^−4^ the recurrence formula was used to compute log fitness, while for smaller values of *μ*_*ss*_ the queuing formula was used.

**Table 5:** Observed mean log fitness for the sexual case under positive selection. Symbols are as described in Table 1. Omitted values required or were expected to require more than 4 core-weeks of CPU time to compute.

| $\mu_{ss}$ | $N$ | $\Gamma_p$ | $s$ | | | |
| --- | --- | --- | --- | --- | --- | --- |
| | | | $10^{-1}$ | $10^{-2}$ | $10^{-3}$ | $10^{-4}$ |
| $10^{-4}$ | $10^5$ | $10^{-3}$ | -0.00712 | -0.00486 | -0.00254 | -0.000749 |
| $10^{-4}$ | $10^5$ | $10^{-2}$ | -0.0694 | -0.0475 | -0.0246 | -0.00716 |
| $10^{-4}$ | $10^5$ | $10^{-1}$ | -0.69 | -0.47 | -0.246 | -0.0702 |
| $10^{-8}$ | $10^5$ | $10^{-3}$ | -1.05 | -0.996 | -1.01 | -1.08 |
| $10^{-8}$ | $10^5$ | $10^{-2}$ | -10.3 | -9.56 | -10.1 | -10.9 |
| $10^{-8}$ | $10^5$ | $10^{-1}$ | -105 | -102 | -100 | - |
| $10^{-9}$ | $10^6$ | $10^{-2}$ | -10.3 | -10.7 | -10.1 | - |
| $10^{-9}$ | $10^5$ | $10^{-3}$ | -10.2 | -9.76 | -9.68 | -10.7 |
| $10^{-9}$ | $10^5$ | $10^{-2}$ | -104 | -95.6 | -98.6 | -109 |
| $10^{-9}$ | $10^5$ | $10^{-1}$ | $-1.01 \times 10^3$ | $-1.02 \times 10^3$ | $-1 \times 10^3$ | - |
| $10^{-9}$ | $10^4$ | $10^{-2}$ | $-1.03 \times 10^3$ | -988 | $-1 \times 10^3$ | -657 |
| $10^{-10}$ | $10^5$ | $10^{-3}$ | -102 | -99 | -97.1 | -108 |
| $10^{-10}$ | $10^5$ | $10^{-2}$ | $-1.01 \times 10^3$ | $-1.01 \times 10^3$ | $-1.02 \times 10^3$ | $-1.09 \times 10^3$ |
| $10^{-10}$ | $10^5$ | $10^{-1}$ | $-1.04 \times 10^4$ | $-1 \times 10^4$ | $-9.96 \times 10^3$ | - |

**Table 6:**
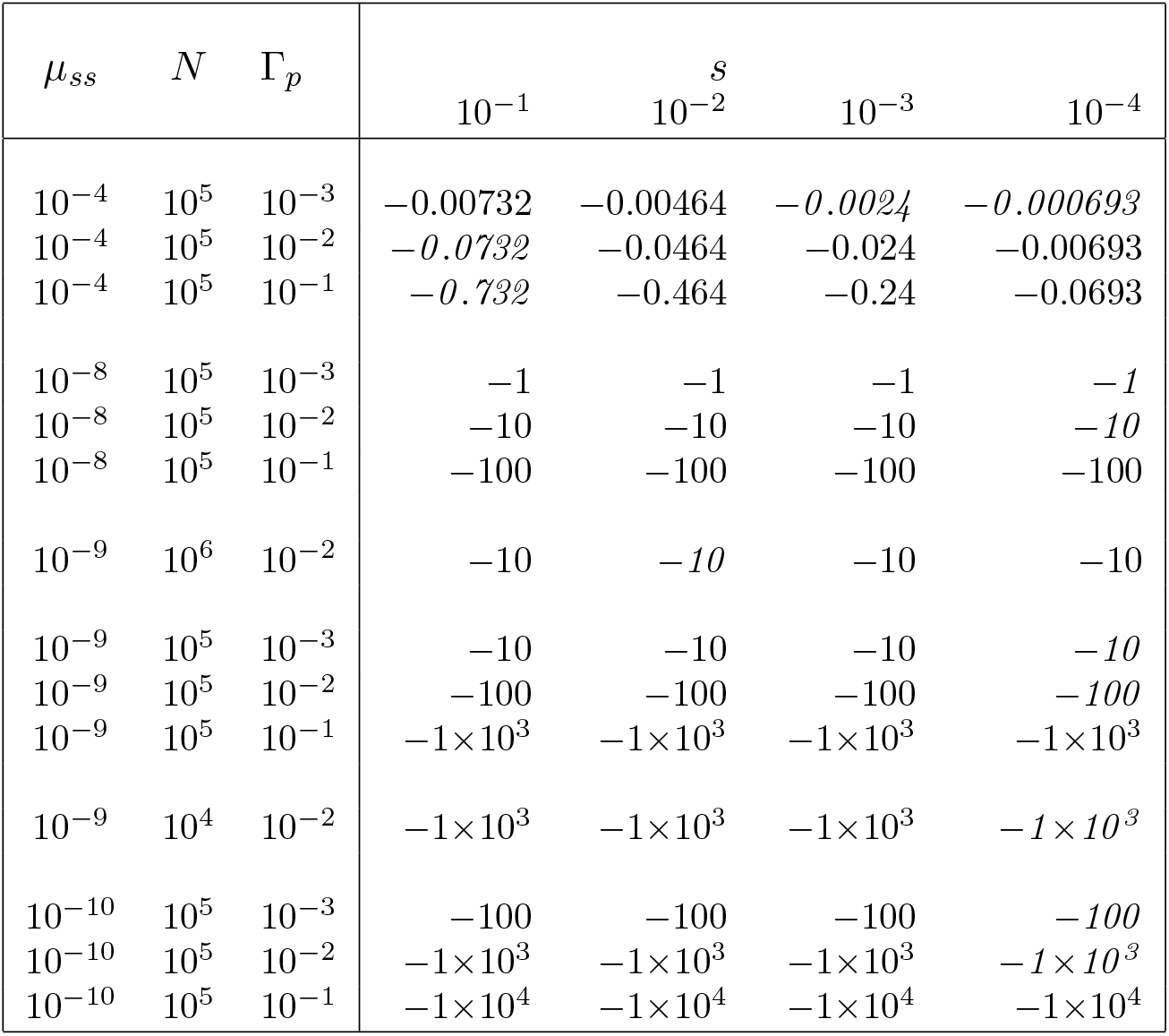
Mean log fitness for the sexual case under positive selection calculated using the larger of the recurrence formula or the queuing formula. Symbols are as described in Table 1.

| $\mu_{ss}$ | $N$ | $\Gamma_p$ | $s$ | | | |
| --- | --- | --- | --- | --- | --- | --- |
| | | | $10^{-1}$ | $10^{-2}$ | $10^{-3}$ | $10^{-4}$ |
| $10^{-4}$ | $10^5$ | $10^{-3}$ | $-0.00732$ | $-0.00464$ | $-0.0024$ | $-0.000693$ |
| $10^{-4}$ | $10^5$ | $10^{-2}$ | $-0.0732$ | $-0.0464$ | $-0.024$ | $-0.00693$ |
| $10^{-4}$ | $10^5$ | $10^{-1}$ | $-0.732$ | $-0.464$ | $-0.24$ | $-0.0693$ |
| $10^{-8}$ | $10^5$ | $10^{-3}$ | $-1$ | $-1$ | $-1$ | $-1$ |
| $10^{-8}$ | $10^5$ | $10^{-2}$ | $-10$ | $-10$ | $-10$ | $-10$ |
| $10^{-8}$ | $10^5$ | $10^{-1}$ | $-100$ | $-100$ | $-100$ | $-100$ |
| $10^{-9}$ | $10^6$ | $10^{-2}$ | $-10$ | $-10$ | $-10$ | $-10$ |
| $10^{-9}$ | $10^5$ | $10^{-3}$ | $-10$ | $-10$ | $-10$ | $-10$ |
| $10^{-9}$ | $10^5$ | $10^{-2}$ | $-100$ | $-100$ | $-100$ | $-100$ |
| $10^{-9}$ | $10^5$ | $10^{-1}$ | $-1 \times 10^3$ | $-1 \times 10^3$ | $-1 \times 10^3$ | $-1 \times 10^3$ |
| $10^{-9}$ | $10^4$ | $10^{-2}$ | $-1 \times 10^3$ | $-1 \times 10^3$ | $-1 \times 10^3$ | $-1 \times 10^3$ |
| $10^{-10}$ | $10^5$ | $10^{-3}$ | $-100$ | $-100$ | $-100$ | $-100$ |
| $10^{-10}$ | $10^5$ | $10^{-2}$ | $-1 \times 10^3$ | $-1 \times 10^3$ | $-1 \times 10^3$ | $-1 \times 10^3$ |
| $10^{-10}$ | $10^5$ | $10^{-1}$ | $-1 \times 10^4$ | $-1 \times 10^4$ | $-1 \times 10^4$ | $-1 \times 10^4$ |

Predictions that differed from observations by more than 2 standard errors are indicated by italics. The two tables are in reasonably close agreement throughout. The major discrepancy being the result for *N* = 10^4^ and *s* = 10^*−*4^, but here *Ns* = 1, which is a boundary case, and none of the formulas in equation 3 are expected to apply. Other values for *s* = 10^*−*4^ appear slightly more negative than predicted. This might because *Ns* is starting to approach 1, and random drift is starting to play a role.

Sexual mean log fitness conjecture: For the sexual model with *μ*_*ss*_ ⪡ *s* ⪡ 1 the mean log fitness calculated using the recurrence formula is given by,

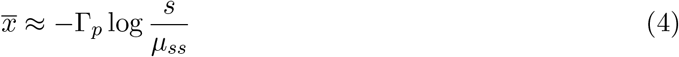

Status: Unproven. Agrees with recurrence formula results shown in Table 6 and additional recurrence formula values calculated but not shown.

### Negative selection and sexual mean log fitness

For the sexual model under negative selection,with 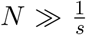, log fitness is approximately given by negating the rate of negative site creation (equation 3). Experiments shown in Table 7 demonstrate this to be correct, but it has not been tested exhaustively.

**Table 7:**
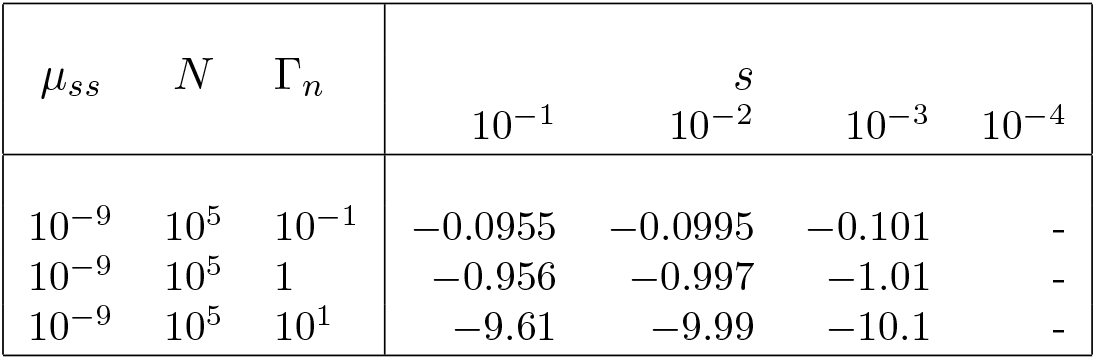
Observed mean log fitness for the sexual case under negative selection. Symbols are as described in Table 1.

| $\mu_{ss}$ | $N$ | $\Gamma_n$ | $s$ | | | |
| --- | --- | --- | --- | --- | --- | --- |
| | | | $10^{-1}$ | $10^{-2}$ | $10^{-3}$ | $10^{-4}$ |
| $10^{-9}$ | $10^5$ | $10^{-1}$ | -0.0955 | -0.0995 | -0.101 | - |
| $10^{-9}$ | $10^5$ | 1 | -0.956 | -0.997 | -1.01 | - |
| $10^{-9}$ | $10^5$ | $10^1$ | -9.61 | -9.99 | -10.1 | - |

### Comparison of the sexual and asexual models

Evaluating the advantage of sex under positive selection could be thought of as an evaluation of the Fisher-Muller hypothesis. Subtracting mean log fitness for positive selection in the asexual case (Table 1) from mean log fitness for positive selection in the sexual case (Table 5) produces Table 8. Omitted sexual values were filled in with the more negative of the recurrence formula or queuing formula (equation 3). This means the recurrence formula was used for *N* = 10^8^, and the queuing formula elsewhere.

**Table 8:** Estimated advantage of sex for positive selection. Observed or predicted mean log fitness for the sexual case less observed mean log fitness in the asexual case. Symbols are as described in Table 1.

| $\mu_{ss}$ | $N$ | $\Gamma_p$ | $s$ | | | |
| --- | --- | --- | --- | --- | --- | --- |
| | | | $10^{-1}$ | $10^{-2}$ | $10^{-3}$ | $10^{-4}$ |
| $10^{-8}$ | $10^5$ | $10^{-4}$ | -0.0443 | -0.0517 | -0.0117 | 0.0686 |
| $10^{-8}$ | $10^5$ | $10^{-3}$ | -0.45 | 0.366 | 6.44 | 2.77 |
| $10^{-8}$ | $10^5$ | $10^{-2}$ | 20.2 | 377 | 274 | 49.9 |
| $10^{-8}$ | $10^5$ | $10^{-1}$ | $2.33 \times 10^4$ | $2.23 \times 10^4$ | $5.35 \times 10^3$ | 682 |
| $10^{-9}$ | $10^8$ | $10^{-2}$ | 6.79 | 862 | $1.5 \times 10^3$ | 464 |
| $10^{-9}$ | $10^7$ | $10^{-2}$ | 19.4 | $1.37 \times 10^3$ | $1.78 \times 10^3$ | 501 |
| $10^{-9}$ | $10^6$ | $10^{-2}$ | 62.1 | $2.17 \times 10^3$ | $2.18 \times 10^3$ | 530 |
| $10^{-9}$ | $10^5$ | $10^{-4}$ | -0.496 | -0.504 | -0.122 | 0.65 |
| $10^{-9}$ | $10^5$ | $10^{-3}$ | -4.39 | 4.43 | 65.9 | 26.9 |
| $10^{-9}$ | $10^5$ | $10^{-2}$ | 205 | $3.76 \times 10^3$ | $2.74 \times 10^3$ | 505 |
| $10^{-9}$ | $10^5$ | $10^{-1}$ | $2.33 \times 10^5$ | $2.23 \times 10^5$ | $5.36 \times 10^4$ | $6.82 \times 10^3$ |
| $10^{-9}$ | $10^4$ | $10^{-2}$ | 545 | $6.91 \times 10^3$ | $2.94 \times 10^3$ | 97.9 |
| $10^{-10}$ | $10^5$ | $10^{-4}$ | -5.29 | -5.23 | -1.16 | 6.24 |
| $10^{-10}$ | $10^5$ | $10^{-3}$ | -46.6 | 44.1 | 635 | 277 |
| $10^{-10}$ | $10^5$ | $10^{-2}$ | $1.75 \times 10^3$ | $3.8 \times 10^4$ | $2.72 \times 10^4$ | $5.1 \times 10^3$ |
| $10^{-10}$ | $10^5$ | $10^{-1}$ | $2.36 \times 10^6$ | $2.22 \times 10^6$ | $5.36 \times 10^5$ | $6.82 \times 10^4$ |

In the real world sex likely incurs additional costs not captured in the model. The increased complexity, time, and resource consumption of meiosis. The cost of finding a mate. And so on. It seems likely that these costs are going to incur no more than a small multiplicative reduction factor to fitness. Or put another way, the log fitness cost is likely to be no more than, say, 2. Everywhere in table 8 that exceeds this value, sexual reproduction can be expected to outperform asexual reproduction. Empirically, apart from for Γ_*p*_ = 10^*−*4^ this is nearly always the case. E.g. for *μ*_*ss*_ = 10^*−*9^, *N* = 10^5^, Γ_*p*_ = 10^*−*2^, and *s* = 10^*−*2^, the log advantage of sex is 3,760. This is a log advantage, raising *e* to this amount produces an approximate advantage of sex of 10^1,633^. Raising *e* to the values in table 8 typically produces large values, so the conclusions are insensitive to the precise value of any additional costs of sex.

It might be expected that sex would always confer some positive or zero advantage. The negative values in table 8 show this is not the case, at least for the model used here. Both the asexual and sexual models involve a weighted sampling with replacement to simulate selection. The act of recombination then involves a second unweighted sampling with replacement to select each parent. No similar second sampling occurs in the asexual case. Sampling each parent in this way for the sexual case reduces the number of organisms that are effectively parents of the next generation. This reduces the odds that a newly occurring mutation will establish itself in the population for the sexual case when compared to the asexual case. The effect of this stands out the most when selection is strong and mutation is weak, In this case no more than one single mutation typically occurs in the population at a time, and sex might initially have been expected to have had no impact. That this additional sexual sampling causes worse performance for the sexual case was verified by picking the worst offender of asexual outperformance, and testing the asexual model with an extra sampling step. The worst offender is *μ*_*ss*_ = 10^*−*10^, *N* = 10^5^, Γ_*p*_ = 10^*−*3^, and *s* = 10^*−*1^. Performing an extra asexual sampling step resulted in a mean log fitness of 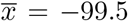, which compares favorably to the predicted value for the sexual case computed using equation 3, 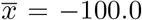. Once the extra sampling is performed, the asexual model no longer significantly outperforms the sexual model.

Evaluating the advantage of sex under negative selection might be thought of as an evaluation of Muller’s ratchet. Subtracting Table 3 from Table 7, or from equation 3 for parameter values that were not simulated, produces Table 9 showing the log advantage of sex under negative selection.

**Table 9:** Estimated advantage of sex for negative selection. Observed or predicted mean log fitness for the sexual case less observed mean log fitness in the asexual case. Symbols are as described in Table 1.

| $\mu_{ss}$ | $N$ | $\Gamma_n$ | $s$ | | | |
| --- | --- | --- | --- | --- | --- | --- |
| | | | $10^{-1}$ | $10^{-2}$ | $10^{-3}$ | $10^{-4}$ |
| $10^{-8}$ | $10^5$ | $10^{-3}$ | 0 | 0 | 0 | 1.14 |
| $10^{-8}$ | $10^5$ | $10^{-2}$ | 0 | 0 | 50.8 | 41.2 |
| $10^{-8}$ | $10^5$ | $10^{-1}$ | -0.00952 | $1.79 \times 10^3$ | $3.23 \times 10^3$ | 659 |
| $10^{-8}$ | $10^5$ | 1 | $7.67 \times 10^4$ | $2.67 \times 10^5$ | $6.06 \times 10^4$ | $8.19 \times 10^3$ |
| $10^{-8}$ | $10^5$ | $10^1$ | $2.46 \times 10^7$ | $5.7 \times 10^6$ | $7.92 \times 10^5$ | $9.08 \times 10^4$ |
| $10^{-9}$ | $10^8$ | 1 | -0.0951 | $1.61 \times 10^6$ | $5.04 \times 10^5$ | $7.53 \times 10^4$ |
| $10^{-9}$ | $10^7$ | 1 | 11.4 | $1.88 \times 10^6$ | $5.32 \times 10^5$ | $7.7 \times 10^4$ |
| $10^{-9}$ | $10^6$ | 1 | $1.28 \times 10^5$ | $2.22 \times 10^6$ | $5.65 \times 10^5$ | $7.93 \times 10^4$ |
| $10^{-9}$ | $10^5$ | $10^{-3}$ | 0 | 0.0752 | 0 | 11.4 |
| $10^{-9}$ | $10^5$ | $10^{-2}$ | 0 | 0 | 504 | 413 |
| $10^{-9}$ | $10^5$ | $10^{-1}$ | 0 | $1.78 \times 10^4$ | $3.24 \times 10^4$ | $6.62 \times 10^3$ |
| $10^{-9}$ | $10^5$ | 1 | $7.65 \times 10^5$ | $2.67 \times 10^6$ | $6.06 \times 10^5$ | $8.19 \times 10^4$ |
| $10^{-9}$ | $10^5$ | $10^1$ | $2.46 \times 10^8$ | $5.7 \times 10^7$ | $7.92 \times 10^6$ | $9.08 \times 10^5$ |
| $10^{-9}$ | $10^4$ | 1 | $2.46 \times 10^6$ | $3.29 \times 10^6$ | $6.63 \times 10^5$ | $8.58 \times 10^4$ |
| $10^{-10}$ | $10^5$ | $10^{-3}$ | 0.499 | 0 | 0 | 113 |
| $10^{-10}$ | $10^5$ | $10^{-2}$ | 0 | 0 | $5.25 \times 10^3$ | $4.14 \times 10^3$ |
| $10^{-10}$ | $10^5$ | $10^{-1}$ | -0.00952 | $1.8 \times 10^5$ | $3.24 \times 10^5$ | $6.55 \times 10^4$ |
| $10^{-10}$ | $10^5$ | 1 | $7.65 \times 10^6$ | $2.67 \times 10^7$ | $6.06 \times 10^6$ | $8.19 \times 10^5$ |
| $10^{-10}$ | $10^5$ | $10^1$ | $2.46 \times 10^9$ | $5.7 \times 10^8$ | $7.92 \times 10^7$ | $9.08 \times 10^6$ |

For negative selection the advantage of sex is usually less than for positive selection for the same parameter values. There is frequently no advantage when *N* is large, Γ_*n*_ is small, and *s* is large.

### Present day prokaryotes

Consider positive selection. Prokaryotes have small *μ*_*ss*_ values, of 10^*−*11^ to 10^*−*9^ mutations per site per generation. If Γ_*p*_ *≤* 10^*−*4^ and *s ≥* 10^*−*3^ there would be an advantage to asexuality for positive selection based on Table 8. To get such a low value for Γ_*p*_ requires a short generation time.

For negative selection under the same *μ*_*ss*_ values, provided Γ_*n*_ *≤* 10^*−*2^ when *s* = 10^*−*2^, there wouldn’t be any disadvantage to asexuality. As mentioned in the introduction, prokaryotes have Γ_*n*_ *≤* 10^*−*2^ mutations per genome per generation. To get this small value for Γ_*n*_ requires a small genome and a small spontaneous mutation rate.

Prokaryotes thrive when the rate of positive and negative site creation per generation is low. They do so by having short generation times, small genomes, and small spontaneous mutation rates.

### Present day eukaryotes

With Γ_*p*_ in the range 10^*−*3^ to 10^*−*2^, based on Table 8, there is typically a large benefit to sex as a result of positive mutation for a wide range of *μ*_*ss*_, *N*, and *s ≤* 10^*−*2^. A new positive mutation with an *s* of 10^*−*1^ is likely to be a very rare event.

With Γ_*n*_ in the range 10^*−*1^ to 10^1^, consulting Table 9, there would be a large advantage to sex except for Γ_*n*_ = 10^*−*1^ and *s* = 10^*−*1^ where the benefits are equivocal.

Thus eukaryotic species today obtain a significant benefit from sex, and do so through both positive and negative selection.

### Early eukaryotic evolution

Sex might make sense for eukaryotes today, but we also need to ask whether it would have made sense for the first eukaryotes, and for that we need to turn to the archaea. Eukaryotes are believed to have evolved from the merger of Asgard archaea and an alphaproteobacterial endosymbiont, possibly in a geothermal environment[4]. The changing nature of geothermal environments may have created a high rate of environmental positive site creation, making it an ideal environment within which for sex to develop.

Spontaneous mutation rates for thermophilic archaea aren’t widely reported, but for the thermophilic *Sulfolobus acidocaldarius* is estimated as 7.8*×*10^*−*10^ mutations per base per generation[5], and for the mesothermophilic *Haloferax volcanii* a rate of 3.2*×*10^*−*10^ has been reported[6]. Thus estimating *μ*_*bs*_ = 5.5*×*10^*−*10^, and *μ*_*ss*_ = 1.8*×*10^*−*10^ might be reasonable. Consequently, for positive selection consulting Table 8, if Γ_*p*_ *≥* 10^*−*3^ sex would have been advantageous for *s* values less than 10^*−*1^. Note that such a large *s* value seems implausible for beneficial mutations.

The known Asgard archaeal genomes range in size from 1.4-5.7M bp, so a typical value for the genome length might be 3.6*×*10^6^ base pairs. For archaea the fraction of the genome that is coding sequence is typically around 0.85[7, Table S1]. If, say, 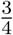 of sites are nonsynonymous sites, then the number of nonsynonymous sites, *A* is 2.3*×*10^6^. Let 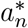 be the fraction of nonsynonymous sites maintained by negative selection, and *n*_*a*_ be the fraction of nonsynonymous sites in the fraction of the genome under the control of negative selection. Then,

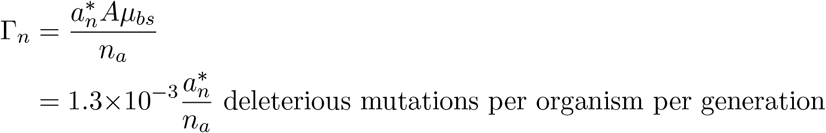

For prokaryotes *n*_*a*_ will be close to 1, making Γ_*n*_ small no matter what the value of 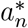. Thus from Table 9 negative selection will have little effect, and so for early eukaryotes Γ_*p*_ would likely be the primary driving force for sex.

Whether Γ_*p*_ exceeded, of the order of 1 site every 1,000 generations, isn’t known for certain. In a changing geothermal environment it seems quite likely, especially considering the many mutations that might be associated with a single simple change in temperature.

Sex frees the genome from the constraints of asexuality. The generation time can be much larger without incurring large costs from positive selection. And the genome can be much larger without incurring high costs associated with negative selection. The possibility of a larger generation time and a larger genome size might explain two important properties of eukaryotes: eukaryotes are both physically much larger and more complex than prokaryotes. The prokaryote typically needs a small generation time and small genome size. The eukaryote can take its time and have a large genome size. The organism can grow larger and be more complex if this offers an evolutionary advantage.

### The mitochondrion as an asexual organism

Comparing human and chimpanzee mtDNA, the non-synonymous substitution rate of protein coding genes is 2*×*10^*−*9^ substitutions per site per year[8]. If we estimate mtDNA contains about 13,500 sites other than synonymous sites, this gives a rate of adaptive mutational opportunity site creation of 2.7*×*10^*−*5^ sites per year, or once every 40,000 years. The time taken for a site to fully fix is approximately 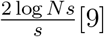, which even for a large population size of 10^6^, a relatively small *s* value of 10^*−*3^ per generation, and a generation time of 25 years, amounts to a full fixation time of 350,000 years. Consequently we can expect the mitochondrial population to contain several haplotypes that all descended from a a single mitochondrial “Eve” that existed perhaps 350,000 years ago. The available evidence suggests Eve actually existed maybe 200,000 years ago[10], implying a larger value for *s* than was imagined.

From the above, for the mitochondrial genome, Γ_*p*_ = 7*×*10^*−*4^ sites per generation. For synonymous sites the mtDNA substitution rate is 3*×*10^*−*8^ per site per year[8]. Thus *μ*_*ss*_ = 2.5*×*10^*−*7^ mutations per generation. Using the estimate of 13,500 sites that are not synonymous, Γ_*n*_ = 1.0*×*10^*−*2^ sites per generation. Extrapolating Table 8, there is likely to be neither a significant cost, nor benefit, to asexuality under positive selection. Recall *s* values for negative selection are likely to be larger than those for positive selection. Based on Table 9, there is likely to be no cost for asexuality as a result of negative selection for *s* = 10^*−*2^ or larger.

The genome size and mutation rate parameters of the mitochondrial genome are such that its asexuality appears to incur little or no cost.

## Discussion

Computer simulation was used to validate the mathematical creation-mutation-selection model.

Although the model shows a strong advantage to sex for a population in certain circumstances, it says nothing about how sex might have evolved. Evolution is not normally viewed as working for the good of the population, only for the good of the gene.

For both sexual and asexual organisms, increasing the spontaneous mutation rate reduces the cost associated with positive selection, but increases the cost associated with negative selection. This is because for finite genomes Γ_*n*_ will scale with the spontaneous mutation rate, *μ*_*ss*_. A balance thus exists between positive and negative selection at which the spontaneous mutation rate is optimal.

Prokaryotes thrive under small rates of environmental change, with short generation times, small genomes, and small spontaneous mutation rates. Sexual organisms thrive under larger rates of environmental change. They are able to have longer generation times and larger rates of negative site creation per genome per generation, and thus larger genomes, and higher spontaneous mutation rates.

For eukaryotes today, and probably the first eukaryotes, the cost of asexuality appears likely to significatly exceed the cost of sex.

## Materials and methods

### Simulation of the asexual model

Simplified pseudo-code for the asexual simulation is shown in Figure 1. See Supplement 1 for the actual simulation code[11].

The implementation was sped up by keeping track of not the number of mutations for each organism, but the number of organisms with each different mutation count. The overall simulation was implemented in Python with the performance critical mutation and selection steps implemented as multi-threaded C code.

Best efforts were made to determine the steady state log fitness mean and variance using a linear regression of the direction in which the mean log fitness was drifting as a function of the initial value for 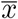. Regression helps overcome the problem of stochastic noise associated with any single parameter run. A variant of equation 1 was used to determine the initial value for 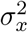 with the initial fitness distribution being log-normally distributed. Regression data points were gathered until the estimated mean log fitness value for zero drift was accurate to within 5% with 95% confidence as determined through bootstrapping. Each regression data point was generated by simulating warm-up time which was not assessed, followed normally by 10,000 generations of measurement time. The measured drift was adjusted for stochastic variation in site creation. The warm-up time lasted until 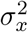 averaged out over preceding, normally 2,000 time-step intervals, had been seen to both increase and decrease.

Once the regression was complete, a simulation was performed starting with a random population with log-normally distributed fitness generated from the estimated steady state log fitness mean and variance computed as described above. Warm-up time was simulated during which no results were collected, followed normally by 10,000 generations of measurement time. The process was usually repeated 100 times to account for long time period fluctuations in 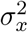, and averages taken. To reduce the compute cost for *N* = 10^8^, just 20 samples of 5,000 generations were generated.

The regression approach fails if 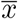 changes significantly from its initial value during the course of the simulation. Consequently, letting Γ represent either Γ_*p*_ or Γ_*n*_, if the regression predicted the expected value of 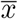 was smaller in magnitude than 50,000Γ log(1+*s*) a direct roll-out was performed. No results were collected until the roll-out appeared stable. The roll-out was considered stable when 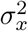 had been observed to both increase and decrease when averaged out over normally 50,000 time-steps, followed by 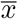 adjusted for stochastic fluctuations in the rate of site creation having been seen to both increase and decrease when averaged over normally 50,000 generations. Once the simulation was stable results were normally collected for 1,000,000 generations divided into 10,000 generation intervals. Direct roll-outs were started from the value of *x* predicted by the partially complete regression.

On account of the low frequency of adaptive site creation for Γ_*p*_ = 10^*−*4^ the number of generations was increased in this case. For the regression warm-up time was averaged using 20,000 generations, followed by 100,000 generations of measurement time, and the final simulation consisted of 20 samples of 100,000 generations. A regression was performed if the expected value for 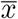 was smaller in magnitude than 500,000Γ_*p*_ log(1 + *s*), with stability assessed over 500,000 generations, and then samples collected for 2,000,000 generations divided into 100,000 generation intervals.

Reported steady state values of 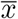 are based on the regression or the roll-out. Estimates of 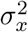 and skewness are based on averages from the final simulation or the roll-out. Standard error values for the roll-outs were adjusted for any correlation between consecutive samples.

### Simulation of the sexual model

Simplified pseudo-code for the sexual simulation is shown in Figure 2. See Supplement 1 for the actual simulation code[11].

**Figure 2:**
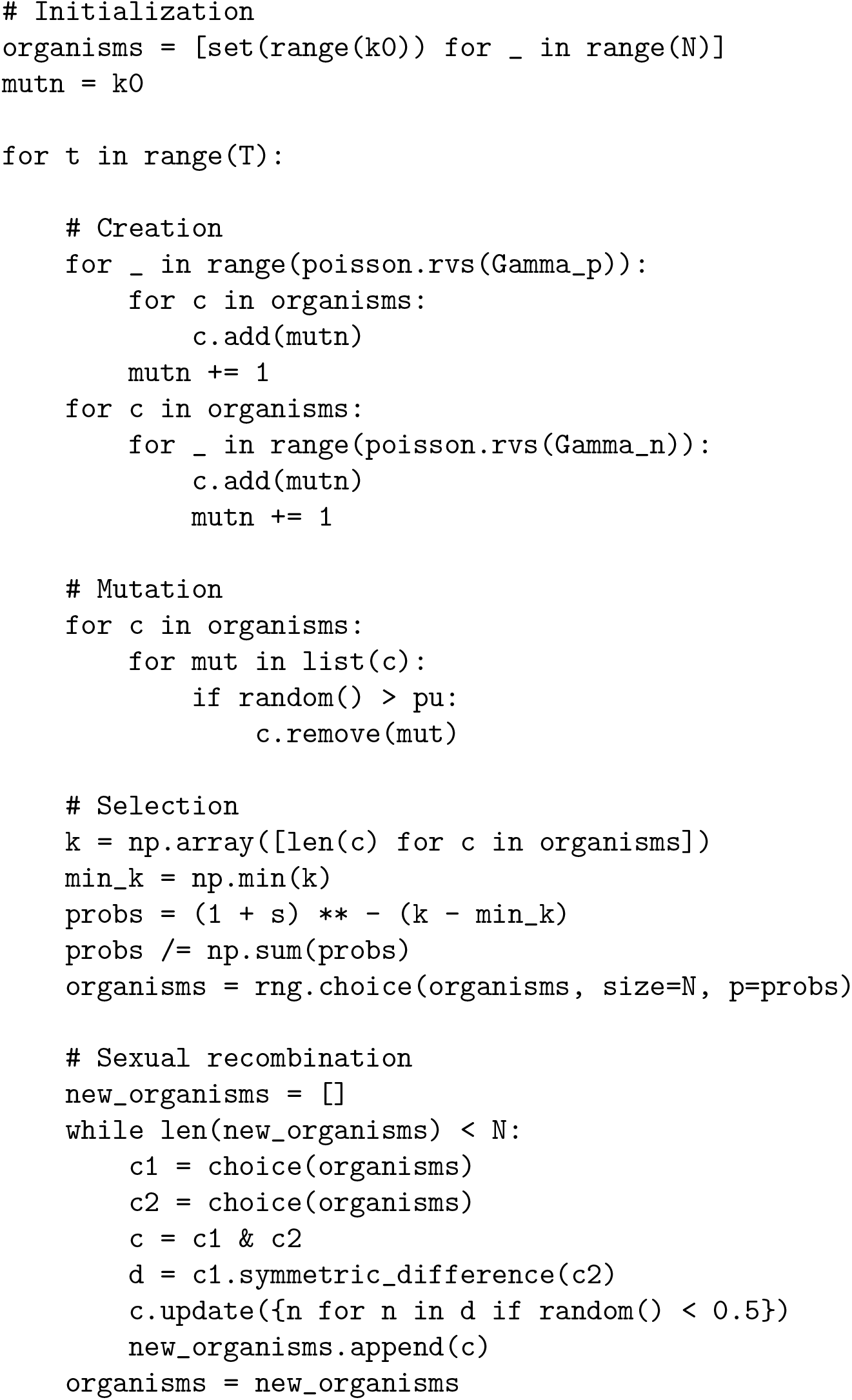
NumPy Python pseudo-code for the sexual simulation.

In the asexual model it is possible to track just the total number of unsatisfied sites in each organism. In the sexual model this is not possible. The identity of each site must be tracked in each organism so that recombination can be correctly computed. Multiplying the number of sites, by the number of organisms, by the number of time-steps, quickly adds up to an excessive amount of computer resources for the simulation. In an attempt to minimize these costs the simulation maintains multiple lists: a global list of sites present in nearly all organisms; a per organism list of exclusions from the global list (for newly satisfied positive sites); and a per organism list of direct sites present in just a few organisms (for new negative sites and sites close to fixation). Periodically these lists were rebalanced: frequently occurring exclusions were replaced by direct list entries and removed from the global list. The simulation was implemented in a mixture of Python, for the non-performance critical regions, and multi-threaded C, for the performance critical regions.

For positive selection two approaches were used for determining the steady state. For expected mean log fitness result values, 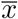, smaller in magnitude than 50,000Γ_*p*_ log(1 + *s*) a direct roll-out was performed starting from the 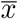 predicted by the recurrence relationship with *σ*_*x*_ = 0. In this case a simulation was judged to have reached a steady state once 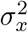 was considered to be stable followed by 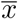 being considered to be stable. 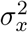 was considered to be stable once its value averaged over the 20,000 preceding time-steps was seen to be both larger and smaller than its value for 20,000 time-steps immediately prior to that. Once 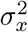 was stable, 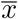 was considered to be stable if both positive and negative changes to its value averaged over the 20,000 preceding time-steps were observed. In making this latter determination 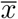 was corrected for stochastic fluctuations in the rate of site creation. An issue with determining the accuracy of the then measured 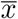 value is that consecutive time-steps will be highly correlated. Results were thus gathered until 100 samples each separated by a sufficiently large stride had an auto-correlation of 0.6 or less. This is equivalent to having about 25 fully independent samples. The accuracy of the resulting 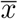 values was then determined by computing their mean and standard error. Finally the simulation was run for an additional 20,000 time-steps and additional results gathered.

For positive selection, running the simulation until a steady state was reached for larger expected values of 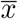 was judged as too time consuming. Instead a regression approach was used. At least 20 starting values for 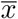 with *σ*_*x*_ = 0 were chosen. For each the system was run until 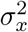 was considered to be stable when averaged, this time, over 200 time-steps. The simulation was then run for 4 times as many time-steps as had occurred up until this point, and the average rate of drift in 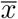 per time-step computed. Over a broad range of starting values, drift was observed to be a linear function of 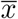. A linear regression was performed of drift as a function of 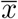, and the value of 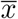 for which drift was expected to be zero computed. Bootstrapping of the linear regression was performed to determine the accuracy of the results. A final simulation at this point of expected zero drift was performed, waiting until 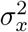 was stable, and then results gathered over 20,000 time-steps.

For negative selection a direct roll-out was always performed. The procedure was the same as for positive selection except that the system was considered stable when both positive and negative changes in value were seen when averaged out over 2,000 time-steps instead of 20,000 time-steps, and the final simulation was only run for 2,000 time-steps instead of 20,000 time-steps to gather additional results.

Reported steady state 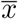 values were based on the mean value of the partially auto-correlated samples for the direct roll-outs, or the zero drift point of the regressions. These values are intended to be accurate to within 5% with 95% confidence. Other attributes of the sexual simulation, such as variance and skewness, were measured based on the final 20,000 time-steps, but are not reported in this paper.

## Supplements

Supplement 1 - Asexual and sexual creation-mutation-selection simulation software. https://doi.org/10.5281/zenodo.8080283

